# Neuronal normalization in monkey MT is an intensity-weighted average

**DOI:** 10.1101/2025.07.21.665985

**Authors:** Chery Cherian, John H. R. Maunsell

**Affiliations:** Department of Neurobiology and Neuroscience Institute, University of Chicago, Chicago, IL 60637, USA

## Abstract

Normalization is a ubiquitous neuronal computation that is important for safeguarding stimulus selectivity. However, normalization strength has been found to vary greatly across neurons. Here, we show that the normalization of responses by neurons in the macaque middle temporal visual area (MT) is profoundly affected by the receptive field responsivity at each stimulus location. An intensity-weighted normalization model, in which intensity is defined as the product of stimulus contrast and a location-specific receptive field weight, explains most of the previously observed variability in normalization across neurons. It furthermore explains systematic changes in the semi-saturation contrast of contrast response functions at different receptive field locations. Finally, intensity-weighted normalization reveals that spontaneous activity can be viewed as unknown excitatory drive that has measurable intensity and contributes to normalization equivalently to experimental stimuli.

## Introduction

Normalization has emerged as a canonical neuronal computation that explains many non-linear responses, not only in visual cortex but across diverse brain regions, sensory modalities, and species, including invertebrates.^1^ It has also been implicated in context-dependent modulation of neuronal representations of cognitive factors, such as spatial attention^2-4^, pairwise noise correlations^5^, working memory^6,7^ and value.^8,9^

Normalization is well-captured by a divisive normalization equation that approximates a contrast-weighted average of the response to the stimuli when presented alone.^10^ This weighted average of responses explains why adding a weakly excitatory stimulus to a strongly excitatory stimulus reduces, rather than increases, a neuron’s response.^10-15^ Importantly, this form of normalization generalizes to conditions involving more than two stimuli within a neuron’s receptive field (RF). For instance, in the middle temporal (MT) area, normalization across all the dots in a random dot kinematograms accounts for the linear relationship between MT neuron firing rates and motion coherence.^16^

Normalization has frequently been suggested to support intensity- or contrast-invariant neuronal sensory representations.^10,14,15,17,18^ By appropriately modulating responses in the presence of multiple stimuli, normalization may play a critical role in preserving neuronal selectivity, as it ensures that no constellation of non-preferred stimuli can elicit a response as strong as a single preferred stimulus. Because natural environments characteristically present multiple stimuli to neuron’s RFs, normalization may be critical for maintaining the fidelity of sensory encoding, particularly in higher stages of sensory processing, where RFs become larger^19^ and response properties become increasing selective.^20^

Despite the wide expression of normalization and its apparent functional significance, some observations suggest that it is not tightly regulated, i.e. there is heterogeneity in the extent neurons normalize their response. While the average normalization across populations of neurons closely approximates a contrast-weighted average of responses, individual neurons vary widely, with some having responses that are much more or far less sub-linear than predicted.^2,3,21^ Normalization for individual neurons can also vary greatly when the RF positions of preferred and non-preferred stimuli are exchanged,^3^ suggesting that the strength of normalizing inputs varies across the RF. The influence of RF position on normalization has not been extensively examined. Many studies of normalization focus on V1, whose small RFs dictate that it is examined with superimposed stimuli that fill the entire RF.^10,13,14,22^ Studies of responses to multiple stimuli in the larger RFs of neurons in extra-striate visual areas have not considered the effects of RF location.^23,24^

To investigate how location within the RF affects normalization, we recorded from MT neurons while presenting pairs of small stimuli at multiple locations spanning their large RFs.^25-27^ We found that MT neuron responses are well predicted by a RF-weighted normalization model, in which the contribution of each stimulus to normalization is weighted by the RF strength at the position where it is presented. When normalization is considered as an intensity-weighted average, in which intensity combines stimulus contrast and RF weight, it appears to be far more precisely regulated in neuronal responses than previously recognized.

## Results

We investigated how normalization is impacted by stimulus location within a RF by recording from 90 isolated MT neurons (43 from monkey A, 47 from monkey M) in two monkeys (*Macaca mulatta*). To systematically test how normalization varies across RF location, we measured neuronal responses to a large set of stimulus configurations specifically designed to vary spatial location and stimulus pairing (Figure 1A).

**Figure 1.**
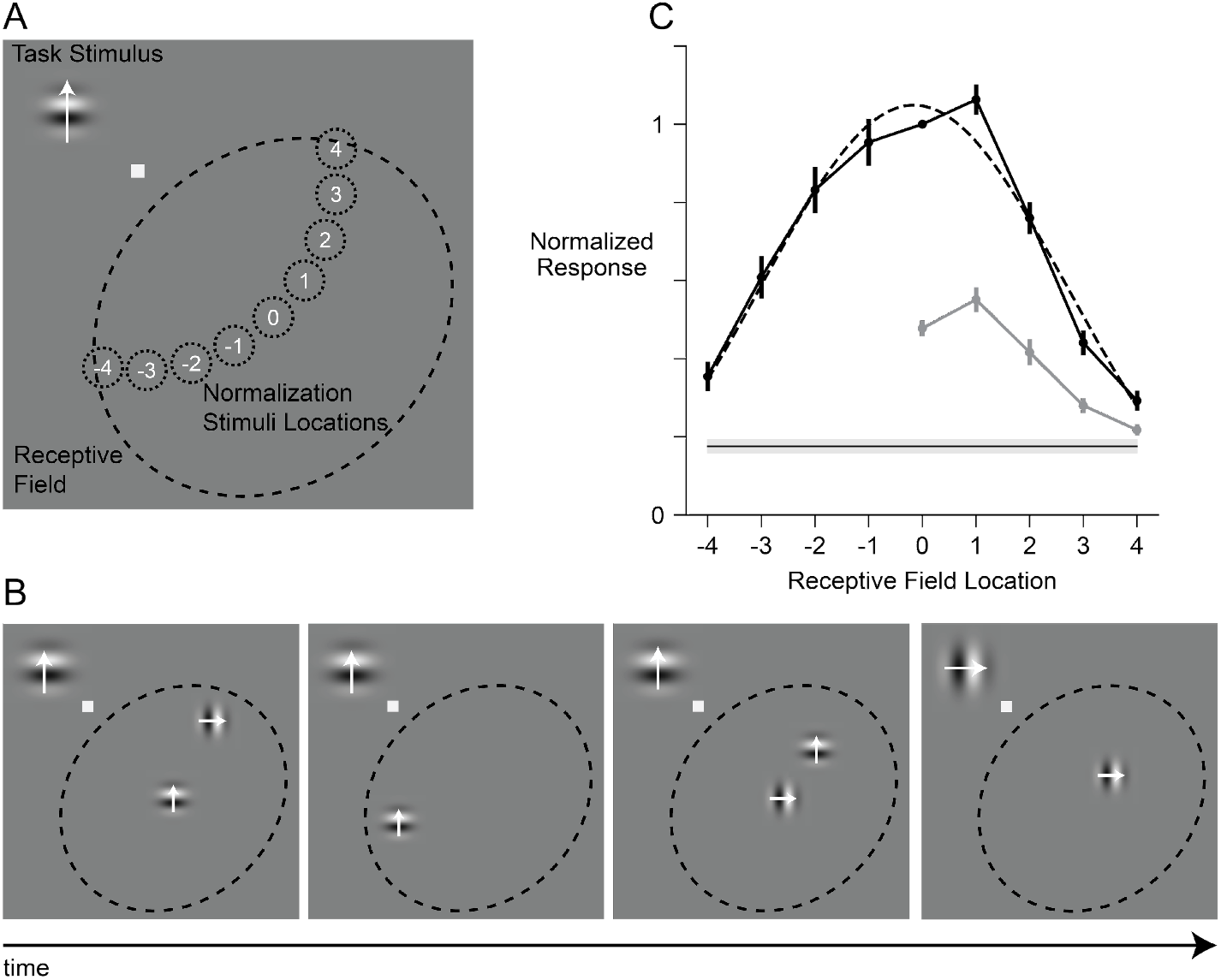
Experimental Design. A) Stimulus layout The monkey performed a change detection task using a central fixation spot and penpheral task-relevant Gabor stimuli located far from the RF Additional Gabor stimuli were briefly flashed within the RF at nine isoeccentric positions to measure the RF profile and characterize normalization. When paired stimuli were presented in the RF. one always appeared at the RF center (location 0) and the other at one of four flanking positions (positions 1-4). RF Gabors drifted in either the neuron’s preferred or a preselected non preferred direction B) Change-detection task. While maintaining fixation, the monkey monitored sequences of task-relevant Gabors to detect a change in orientation. On each trial. 0.1. or 2 Gabor stimuli we re flashed within the RF, independent of the task stimuli C) Average RF profile of MT neurons The mean normalized spatial response profile (n = 90 neurons) was approximately Gaussian. Responses to preferred-direction Gabors are shown in black (solid line, data; dashed line. Gaussian fit). Responses to non preferred Gabors are shown in light gray. The thin horizontal line indicates the mean spontaneous rate. Error bars represent ± SEM.

### MT Receptive Field Profiles

During recordings, the monkeys did a change detection task (Figure 1B). Each trial began when a fixation spot appeared on a video monitor. Shortly after the animal fixated on this spot, a sequence of Gabor stimuli appeared in the left visual hemifield (contralateral to the recorded neurons). Each Gabor was randomly selected to drift in either direction of its drift axis for 200 ms and was separated from other Gabors by randomly selected periods of 200-400 ms. Monkeys were rewarded with juice if they correctly detected the randomly timed appearance of a Gabor with a different drift orientation.

While the animal attended to the task stimuli, we presented sequences of single or paired full-contrast Gabors within the RF of a recorded neuron to measure normalization. The direction and speed of the RF Gabors were matched to the preferences of the cell being recorded. To measure the RF profile, single Gabors drifting in the preferred direction were presented at each of nine iso-eccentric locations spanning the full width of the RF. To test how spatial arrangement influences normalization, pairs of Gabors were presented, with one always placed at the RF center (Figure 1A, location 0) and the other positioned at one of four increasingly offset locations (Figure 1A, locations 1-4) within the same half of the RF. Normalization measurements were restricted to one side of the RF on the assumption that RF effects were spatially symmetric. During paired presentations, the drift direction was either the neuron’s preferred direction, or a fixed non-preferred direction, which was selected to evoke a reliable but weaker response. Each RF stimulus was randomly and independently assigned to have a preferred or non-preferred direction. RF Gabors were presented for 250 ms and were separated from others by 200 ms. Because the RF stimuli were never relevant to the behavioral task, attention-related modulations of neuronal responses were minimized.^2,28-30^

The average RF activity profile of the recorded MT neurons was well-approximated by a Gaussian function (Figure 1C, dotted line; R^2^=0.97). For individual neurons, Gaussian fits to their RF profiles yielded a median explained variance of 92% (interquartile range: 83%-96%) A one-sided mapping of responses using a non-preferred direction revealed a scaled version of the response to the preferred stimulus (Figure 1C, gray points).

Importantly, responses to the selected direction remained above the spontaneous rate of firing at all offset locations, confirming that all tested locations laid within the spatial extent of the RF and thus contributed meaningfully to the excitatory drive (Figure 1C, thin black line). Gaussian RF profiles are the norm for visual neurons in the retina,^31^ LGN,^32^ V1,^33^ and extrastriate areas,^34^ including MT.^24,35,36^

### RF Weighted Normalization

Neuronal responses to paired stimuli are often modelled using the divisive normalization equation:^10^

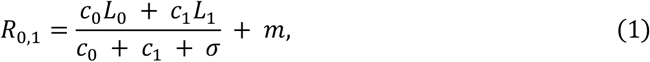

where *c*_0_ and *c*_1_ are the contrasts of two different stimuli (at two distinct sites), *L*_0_ and *L*_1_ are the linear responses to the individual stimuli (e.g., preferred (*L*_*P*_) or non-preferred (*L*_*n*_) at either site), *σ* is a stabilization constant, and *m* is the spontaneous rate. Without *σ* and *m*, the equation is simply a contrast weighted average of the responses to the two stimuli when presented alone. Notably, *σ* is a mathematical convenience to prevent division by zero, but unlike the other terms, which correspond to measurable physical quantities, it lacks a clear biological interpretation, an issue we revisit later in the manuscript. Although, this equation can be expanded to accommodate any number of simultaneously presented stimuli,^16^ here we will consider only pairs of preferred and non-preferred stimuli.

We found that this simple contrast-weighted model did a poor job of capturing response normalization for pairs of stimuli presented across a wide range of RF locations. It typically provided an excellent fit to responses to pairs of stimuli that were both near the RF center, as for the example cell in Figure 2A. The preferred stimulus alone (black line, location 0) produced a much stronger response than the non-preferred stimulus at location 1 (gray line). When these two were presented together (green line) the response closely approximated the average of the individual responses (dashed green line), as has been widely reported.^11,14^ However, when the non-preferred stimulus appeared at a less responsive location in the same cell’s RF (location 2, Figure 2B), the paired response was much less reduced from that to the preferred stimulus alone, similar to weakly normalizing responses previously described in MT.^3,37^

**Figure 2:**
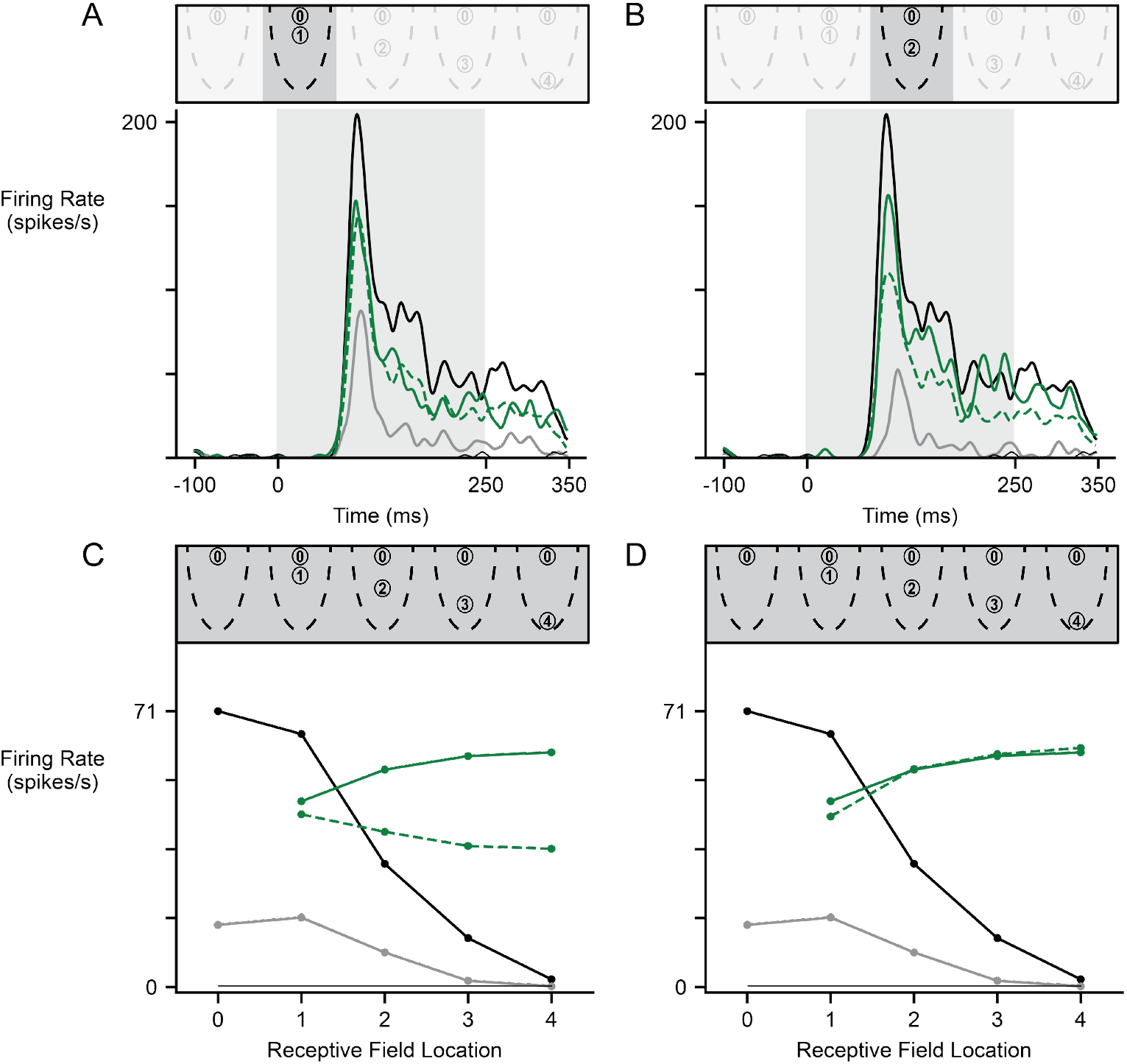
Example neuron responses fit with and without RF-weighted normalization. A) PSTH for an example neuron. Gray shaded region is the stimulus on period. When preferred and non-preferred stimuli are presented together (solid green), the response approximates the average of the responses to each alone (preferred location 0: black line; non-preferred location 1: gray line) The dashed green line shows the average of the two component responses Thin black line shows the spontaneous rate. The schematic above illustrates the stimulus locations within the RF B) Same euron and conventions as in A), but with weaker normalization when the non-preferred stimulus was in location 2. The paired response (green line) deviates from the simple average prediction (dashed green line). C) Responses from the same neuron across a full range of non-preferred stimulus locations. Responses are shown to the preferred direction alone (black), the non-preferred direction alone (gray), and the preferred direction at ocation 0 paired with non-preferred direction at locations 1-4 (solid green). The dashed green line is the average of preferred direction and the non-preferred direction at the offset positions. Thin black line indicates the spontaneous rate. D) Same data as in C) but with the prediction (dashed green line) derived from the RF-weighted model instead of a simple average RF-weighting accomodates the diminishing effect of the non-preferred stimulus as it approached the edge of the RF.

The limitations of Equation 1 become more apparent when applied to the full set of stimulus configurations. Figure 2C illustrates this with responses from the same neuron to a subset of stimulus configurations. Responses to an individual preferred (black line) or individual non-preferred stimulus (grey line) fall off as stimuli approach the RF boundary. Equation 1 specifies that the response to paired stimuli (preferred at location 0 with non-preferred at varying locations) should approximate the average of the component responses (dashed green line). That is, as the non-preferred stimulus shifts toward the RF edge and its individual response decreases, the paired response is expected to decline. However, the measured response instead increases as the non-preferred stimulus approaches the RF boundary (solid green line). This discrepancy arises because Equation 1 lacks spatial specificity – it assumes uniform weighting (i.e., identical *L*_*P*_ and *L*_*N*_) across the RF and therefore cannot accommodate location-dependent contributions.

To account for the varying responses to preferred and non-preferred stimuli at different locations, we extended Equation 1 to include a RF weight term:

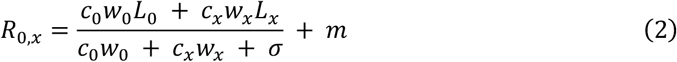

In Equation 2, the *w* term is effectively the relative strength of response to a stimulus presented at different locations. We fix the weight at the RF center (location 0) at 1, while the weights at other locations are treated as free parameters. The subscript *x* signifies the spatial location of offset stimuli, of which there were 4 in our measurements (*x* = 1, 2, 3, 4). Figure 2D shows the responses from the same neuron fit with Equation 2, showing that it captures the reduced influence of stimuli in offset RF locations.

### RF Weighted Normalization Precisely Predicts Responses

Figure 3 shows the MT population average responses for all stimulus conditions fit by the RF weighted normalization model (Equation 2). Twenty-seven response measures are fit based on 8 free parameters. The model captured the data well, explaining 99% of the variance at the population level. To assess whether the RF-weighted normalization model (8 parameters) provided a significantly better fit than the classic normalization model (4 parameters), we compared their Akaike Information Criterion (AIC) scores, which quantify model performance while penalizing for additional parameters. The RF-weighted model yielded lower AIC values for every neuron in the population, with a median ΔAIC of 44.06. Since a ΔAIC > 10 is generally considered strong evidence in favor of one model over another, this large median difference indicates that the RF-weighted model provided substantially better fits, even after accounting for its greater complexity. A Wilcoxon signed-rank test confirmed that this improvement was statistically significant (p<10^−15^).

**Figure 3.**
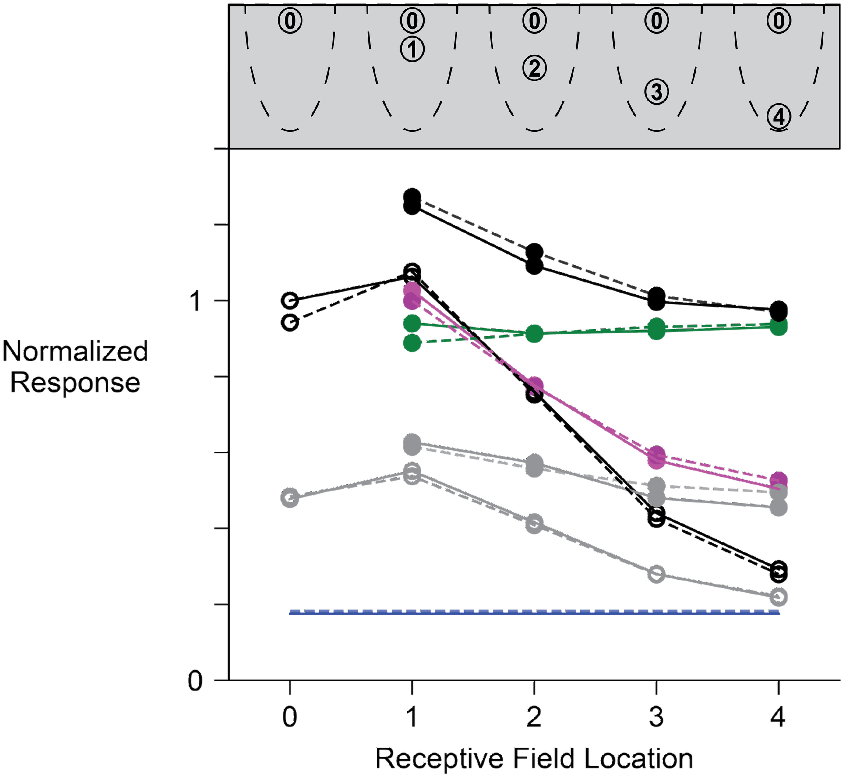
RF-weighting is robustly expressed in normalization across the MT population. Population-averaged normalized responses (solid lines) and corresponding model predictions (dashed lines) are shown (or all stimulus conditions (n = 90 neurons). The RF-weighted normalization model closely tracks the measured responses across stimulus configurations. Model predictions are color-matched to the c rresponding measured data Open circles represent responses to single stimuli; filled circles represent paired stimulus responses Black lines (open circles): Preferred stimulus alone at each location. Gray lines (open circles). Non-preferred stimulus atone at each location. Green lines (filled circles): Preferred stimulus at location 0 paired with a non preferred stimulus at an offset location Black lines (filled circles): Preferred stimulus at location 0 paired with another preferred stimulus at an offset location Magenta lines (filled circles) Non-preferred stimulus at location 0 paired with a preferred stimulus at an offset location. Gray lines (filled circles): Non-preferred stimulus at location 0 paired with another non-preferred stimulus at an offset location Blue line: Mean spontaneous activity

Notably, the RF-weighted normalization model accurately captured several counterintuitive response patterns observed in the data. For example, in the *P*_0_*N*_*x*_condition, where a preferred stimulus appeared at the RF center and a non-preferred stimulus was placed at different offset locations, the model predicted the relatively flat response across offset locations, despite the reduction in the non-preferred stimulus as it approached the boundary (Figure 3; measured: green solid line; predicted: green dashed line). Similarly, in the *N*_0_*P*_*x*_ condition, where a non-preferred stimulus was at the center and a preferred stimulus was placed at different offset locations, the model correctly captured the response rising above the preferred stimulus as the preferred stimulus approached the RF boundary (Figure 3; measured: magenta solid line; predicted: magenta solid line). Equation 2 also accounted for a subtle increase in response to the *P*_0_*P*_*x*_ condition, where two preferred stimuli were presented, one at the center and one at different offset locations, specifically at the nearest offset location (location 1), where the paired response slightly exceeded that of the preferred stimulus alone at the center (Figure 3; measured: black solid lines with filled circles; predicted: black dashed lines with filled circles). This bump in response arises from the influence of the *σ* term in Equation 3. This *σ* term acts as a semi-saturation constant, determining where on the input axis the response begins to plateau. A higher *σ* shifts the semi-saturation point to the right, effectively, dampening the denominator’s growth and allowing additional inputs (like the offset preferred stimulus) to produce modest increases in response. Thus, when responses are not yet saturated, the model can predict slight bumps in activity. In fact, this also explains why during the *P*_0_*N*_*x*_ and *N*_0_*P*_*x*_ condition at the first offset, the paired responses are slightly higher than the pure weighted average.

Across all individual MT neurons, Equation 2 explained a median 94% (90%-97% IQR) of response variance, or 96% (92%-98% IQR) of explainable variance (see Methods). An alternative model that did not apply RF weighting in the denominator performed less well. Across individual neurons, the full RF-weighted model was superior (median explained variance: 94% vs. 92%, p<10^−7^, W=3415; Wilcoxon signed-rank test). Thus, incorporating the RF weighting into the denominator significantly improves the overall fit to neuronal normalization (with the same number of free parameters). In summary, we found that including a RF weight term in both the stimulus drive (numerator) and the normalization (denominator) greatly improves predictions of normalized neuronal responses when stimuli appear at different RF locations.

### RF Weighted Normalization Predicts Changes in Semi-saturating Contrast

One of the notable features of the original divisive normalization equation is that it accounts for the sigmoidal shape of contrast response functions.^10^ When a single stimulus is presented, the equation reduces to:

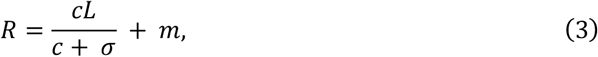

which is a sigmoid function saturating at *m* (lower bound) and *L* + *m* (upper bound). The response is halfway between the saturating values when *c* = *σ*, which is why *σ* is often called the semi-saturation contrast.

However, the RF weight term in RF-weighted normalization affects the contrast at which responses are semi-saturated. For a single stimulus, RF-weighted normalization reduces to:

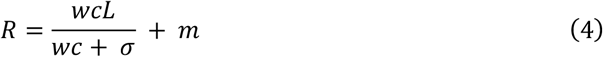

This means the response will be semi-saturated when *c* = *σ*/*w*. We can view the product of RF weight and contrast as setting the effective intensity of a stimulus, in which case *σ* becomes a semi-saturation intensity rather than the semi-saturating contrast. This implies that when RF weight gets larger, a smaller contrast is needed to reach a semi-saturating response.

To explore whether neurons show this behavior, in a separate set of experiments we measured contrast response functions at strong and weak RF locations for 26 isolated MT neurons (20 from monkey A; 6 from monkey M). One location was near the center of the RF while the other was near the RF boundary. At both locations we tested six contrast levels and two directions while the monkey performed the previously described task (Figure 1B). Normalized population averages for each combination of direction and RF location were separately fit with a Naka-Rushton function.

The preferred direction (Figure 4, filled symbols) produced stronger responses than the non-preferred direction (open symbols) whether stimuli were located in the central RF (solid lines) or offset RF (dashed lines). As expected, contrast at semi-saturation at both locations was unaffected by stimulus direction (changing *L* in Eq. 3 or 4). However, at the offset RF location the semi-saturating contrasts for both directions were nearly 50% higher than those at the RF center (9.1% and 9.1% vs 6.4% and 6.6%), consistent with the idea that more contrast is required to reach semi-saturating intensity when the RF weight is weaker (Equation 4).

**Figure 4.**
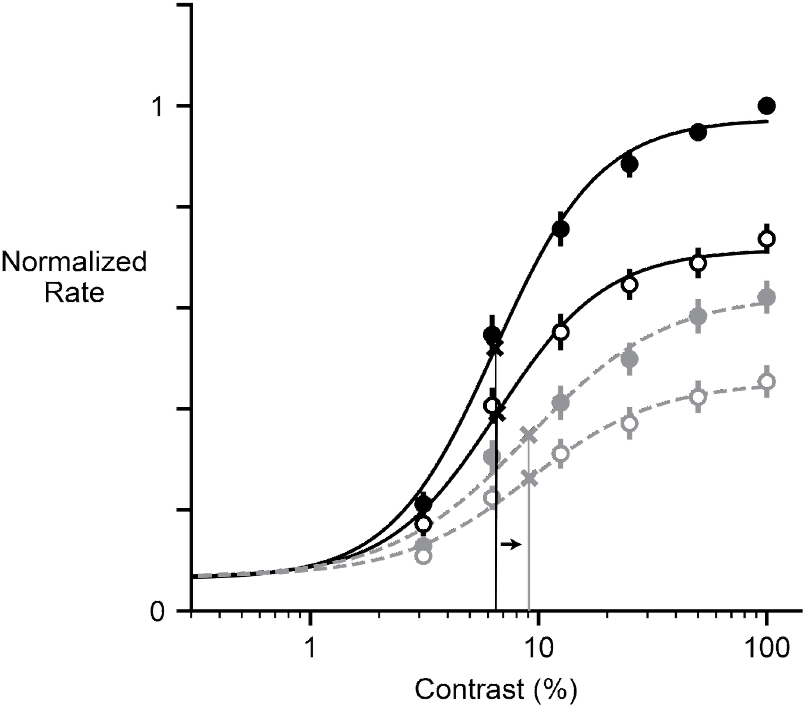
RF-weighted normalization predicts spatial shifts in contrast sensitivity across the RF. Population-avciagcd. normalized contrast response functions are shown for stimuli presented at two RF locations (center and edge) and in two directions (preferred and non-preferred) (n = 26 neurons) Stimuli at the RF center (solid black lines) produced similar contrast sensitivity for both directions (preferred: c_50_ = 6 44; non-prefcrred. c_50_ = 6.58). Stimuli at the RF edge (dashed gray lines) showed reduced contrast sensitivity, with higher c_50_ values (preferred: c_50_ = 9.05: non-preferred: c_50_ = 9.10). Filled circles represent responses to preferred-direction stimuli; open circles represent responses to non-preferred-direction stimuli Black arrow indicates the shift in c_50_ from center to periphery. X marks the denote the half-maximum response for each curve. Thin vertical lines (black for center, gray for edge) indicate the contrast levels corresponding to the half maximum responses Error bars show ±1 SEM

This result was borne out in the fits for individual neurons. There was no significant difference between the distributions of semi-saturating contrasts for preferred versus non-preferred directions for individual neuron responses at either RF location (RF center, p>0.25, U=376.0; RF periphery p>0.34, U=361.0; Mann-Whitney one tailed U test, Bonferroni corrected). However, the semi-saturating contrast were significantly larger in the peripheral RF than in the central RF (preferred direction p<0.025, U=448.0; non-preferred direction p<0.025, U=463.0, Mann-Whitney one tailed U test, Bonferroni corrected).

### Normalization as an Intensity-Weighted Average

Viewing *σ* as an intensity allows for a biologically-interpretable reformulation of the divisive normalization equation that makes it biologically interpretable. In the standard divisive normalization (Equation 2), the contrast (*c*), tuning (*L*) and spontaneous activity (*m*) terms are grounded in physical inputs and neuronal activity. In contrast, *σ* does not correspond to any specific physical stimulus or identifiable biological mechanism.

Instead, it serves as an abstract parameter that sets the scale of normalization. While *σ* can be estimated from data, it should be interpreted as a computational construct that reflects how a neuron manages gain, rather than as a direct proxy for any particular neuronal or physical quantity.

RF-weighted normalization suggests an alternative formulation of normalization that eliminates this abstract term. As noted above, RF weight can be viewed as working with stimulus contrast to define the effective intensity of each stimulus. We can replace each occurrence of *wc* with a corresponding intensity, *i*. We can further propose that neurons do not exhibit truly spontaneous activity; rather, their baseline firing reflects an unidentified excitatory drive that, like stimulus-driven activity, contributes to both the excitatory input and to normalization. In this view, *σ* becomes the effective intensity of those uncharacterized inputs. Although those inputs might arise from top-down sources for which contrast and RF location have no meaning, their intensity can nevertheless be measured based on how they normalize responses to visual inputs. This reformulation of neuronal normalization is expressed by the equation:

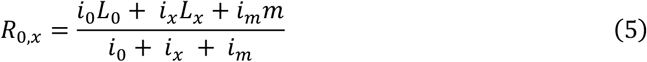

In Equation 5, so-called spontaneous activity is no longer an added offset. Instead, the linear contribution of the unidentified excitatory drive can now be placed in the numerator, weighted by its intensity. In the absence of excitatory stimuli (*i*_0_ = *i*_*x*_ = 0) Equation 5 simplifies to leave only *R* = *m*, which is the baseline firing rate driven by unknown excitatory inputs. But now all inputs, visual or unidentified, act equivalently – each normalizes all the rest – and the abstract *σ* term is no longer needed. Moreover, this reformulation transforms normalization from approximating an contrast-weighted average (Equation 1) to being precisely an intensity weighted average.

Equation 5 arguably maps the normalization equation more closely to physical stimuli and biological reality; however it is not mathematically equivalent to Equation 2, in which the *m* term is applied additively. To compare their performance, we fit responses from each of our neurons to Equations 2 and 5. The two equations exhibited virtually identical performance, with the mean ratio of explained variance deviating from unity by less than 10^−9^ (SEM 10^−10^; two-tailed Wilcoxon signed-rank test W=1919.0, p=0.61). This extraordinary consistent performance in part reflects the fact that undriven activity operates largely like an arithmetic offset, which has no effect on response variance.

While both models perform equally well, Equation 5 carries distinct benefits over Equation 2: it replaces an abstract parameter with a biologically interpretable term and generalizes normalization to the full spectrum of a neuron’s excitatory drive by recasting it as a simple intensity-weighted average.

## Discussion

### Intensity-Weighted Normalization

The findings presented here extend our understanding of neuronal response normalization by revealing that normalization is greatly affected by RF weight. MT responses to a given stimulus decrease progressively as that stimulus is moved from the RF center toward its edge, and those changes in responsivity are accompanied by proportionately weaker contributions to normalization. Incorporating this RF weighting into the normalization model was essential for precisely predicting neuronal responses. RF weight and stimulus contrast combine multiplicatively in the normalization equation to become an intensity term, which captures two distinct factors that modulate the influences of a given stimulus on a neuron’s response. With RF weight included, normalization is transformed from an approximation of a contrast-weighted average into an exact intensity-weighted average.

Recasting normalization as an intensity-weighted average simplifies the computational framework and offers a clearer interpretation of its components. In this view, all inputs, whether they be from stimuli under experimental control or those that create baseline activity, contribute both to driving spikes and to normalization according to their relative intensities. The *σ* term In the classic normalization model (Equation 1) originates from the semi-saturation constant of the Naka-Rushton function, which itself is a form of the Michaelis-Menten equation.^38^ In those contexts, *σ* is a mathematical parameter that shapes the sigmoidal form of the response function but lacks a clear biological interpretation or physical embodiment. Intensity weighted normalization allows for a more neurobiologically plausible alternative: by reinterpreting *σ* as *i*_*m*_, it instead reflects the intensity of unidentified inputs to the neuron – such as local contextual influences, or top-down feedback from higher-order areas (Equation 5). In this way *i*_*m*_ appears as an additional static input that contributes to normalization, in the same way that all experimentally controlled inputs with explicit contrast and RF weight terms. This reframing provides a conceptually simpler and more interpretable framework for evaluating the role of this term in generating normalized neuronal responses.

While the intensity terms associated with visual stimuli are readily decomposed into contrast and RF weight, that is not true for the intensity of undriven activity, *i*_*m*_. Although baseline activity is driven by excitatory inputs, those inputs are likely to derive from long-range inputs which have no RF weight, or top-down or modulatory inputs with no relationship to either the RF weight or visual contrast. While not decomposable, the intensity of those inputs can nevertheless be measured in units directly commensurate with those for the intensities of visual stimuli by assessing their influence in normalizing responses to visual stimuli that are under experimental control. In this way, normalization provides an approach to evaluate the relative strength of a neuron’s ongoing unknown inputs relative to that of its preferred stimuli. For example, it would in principle be possible to show that a switch between experimental contexts altered the strength of unidentified inputs (*i*_*m*_) but had no effect on the baseline spike rate (*m*).

The ability to treat intensity independently of RF weight or contrast might be useful for approaching normalization in regions beyond sensory cortices that have no direct analog for these terms. In regions supporting more cognitive representations, intensity might apply to non-spatial dimensions such as task relevance, working memory content, or internal state. For example, in the lateral intraparietal cortex (LIP) neuronal signals that represent the value of different choices^8,9^ are normalized as the value of other options change.

Our results align well with a study by Ferrera and Lisberger^39^ that found that the responses of MT neurons to a preferred stimulus near the RF center was little affected by another stimulus near the RF edge. However, previous studies of normalization have not emphasized the importance of RF weight. Many measurements have involved V1, where small receptive fields make spatial offset stimulus presentations difficult, and superimposed stimuli, such as cross-orientation gratings, are commonly used,^1^ masking the effects of RF weighting. Britten and Heuer^24^ presented pairs of stimuli at different locations in MT RFs, but used two preferred stimuli rather than mixtures of preferred and non-preferred stimuli. Consequently, responses appeared to be winner-take-all (as in the solid black or gray symbols in Figure 3), and they concluded that moving a second stimulus towards the RF boundary has little impact on neuronal response.

Precise RF measurements have shown that some MT neurons have RFs with profiles that deviate significantly from a 2D Gaussian,^36^ which might arise from spatially inhomogeneities in the ratio of excitatory and inhibitory inputs.^40^ Because RF-weighted normalization can assign weights based on measured responses, it can accommodate such irregular RF structure. It might be extended to include integrals of detailed stimulus and RF profiles.

### Intensity-Weighted Normalization and Variance in Normalization Strength

Previous work from our lab has reported that normalization strength can vary markedly between MT neurons.^2,3^ In that work, recordings were done with pairs of stimuli were presented at two fixed RF locations that were selected to have similar responses (median stimulus separation 4.2°). RF-weighted normalization suggests that any unaddressed mismatch in RF weight between two sites will affect the apparent strength of normalization, meaning that much of the reported variability might arise from differences in RF weight between the selected sites. In Ni and Maunsell,^3^ normalization was modelled using an equation that allowed suppression to differ between stimulus locations but did not link those changes to differences in excitatory drive. When we used that equation to fit stimulus pairs from our dataset from locations that had the most similar RF weight (median stimulus separation 4.6°), the median explained variance was 93%. In contrast, the RF-weighted normalization model captured significantly more variance under the same conditions (median: 97% vs. 93%; p<10^−12^, W=3761; Wilcoxon signed-rank test) with the same number of parameters.

Previous work used a normalization modulation index (NMI) to measure the strength of normalization.^2,3^ This metric compares a neuron’s response to two stimuli with both the linear sum and the average of the individual responses, capturing the degree to which responses deviate from linear summation. These studies reported substantial variability in normalization across and within neurons. More recently, modeling work has proposed that this heterogeneity may actually be beneficial, as it enhances the population’s capacity to encode visual information.^41^ However, our findings offer a different interpretation. Rather than reflecting true differences in normalization strength, the apparent variability may arise from imbalances in the spatial drive of the stimuli. Our results suggest that MT neurons apply a consistent normalization computation, an intensity-weighted average of the inputs, rather than a simple contrast-weighted average. Classic normalization mechanisms serve to preserve neuronal selectivity by preventing multiple non-preferred stimuli from eliciting the same response as to a single preferred stimulus. RF weighted normalization or intensity weighted normalization extends this principle by preserving both direction and spatial selectivity, ensuring that a neuron fires at its maximum rate only under optimal conditions. From this perspective, the variability in suppression observed across the population may not reflect differences in normalization mechanisms per se, but rather differences in how strongly the second stimulus engages the RF. When the second stimulus falls within a more responsive RF region, it contributes more strongly to the normalization pool and modulates the response more effectively – creating population-wide variability that may enhance the encoding capacity of the population.

### Influence of RF Weighting on Semi-Saturation Contrast

A notable success of the RF weighted normalization model is its ability to predict changes in the half-maximum of the neuron’s contrast response function depending on where within the RF the stimulus is presented. Contrast response functions measured in locations with higher RF weights have lower semi-saturation contrasts, while those at weaker locations have higher semi-saturation contrasts. These predictions are consistent with findings by Heuer and Britten,^42^ who observed spatial variation in contrast response functions across the RF. However, they focused their analysis on the changes in response amplitude, as that explained most of the variance and as a result downplayed changes in the semi-saturation contrast. In contrast, our results show that both amplitude and sensitivity vary significantly across the RF, with sensitivity shifts reaching statistical significance. This finding aligns with prior reports of reduced contrast response function sensitivity at the periphery of classical RFs and into the surround in V1 and MT,^43,44^ although direct comparisons are limited because our stimuli were restricted to the classical RF edge.

Systematic differences in semi-saturating contrast have been seen previously in MT receptive fields. Sclar and his colleagues^45^ measured contrast responses functions from individual MT neurons while varying the radius of a drifting grating patch. They found that semi-saturating contrast decreased as patch size increased. This is expected based on RF-weighted normalization. A step in patch radius can be viewed as adding another stimulus to the RF, changing the denominator in Eq. 4 from *w*_0_*c* + *σ* to *w*_0_*c* +*w*_1_*c* + *σ*, thereby reducing the semi-saturating contrast from *c* = *σ*⁄*w*_0_ to *c* = *σ*/(*w*_0_ +*w*_1_). If we accept that MT RFs have a Gaussian weighting profile (Figure 1), the rate at which semi-saturating contrast decreases with patch radius will not be constant, but for a circular patch with a radius of ∼ 1 *SD* of the RF Gaussian, semi-saturation contrast will be proportional to 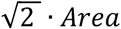, which is the value reported by Sclar and colleagues.^45^

We note that Equation 4 does not capture all the differences in responses. In particular, there is an apparent change in the R_max_ for both stimulus directions in going from the center to the periphery (Figure 4). Equation 4 does not produce a change in R_max_ with changes in RF weight. However, the change we observe in both R_max_ and the semi-saturation contrast is consistent with earlier work that measured contrast response functions at different locations within the RF.^42^ Equation 4 also suggests that the approximately two-fold change in RF weighting seen in Figure 4 should be accompanied by a proportional increase in the semi-saturating contrast. The measured change is appreciably less than this. Nevertheless, the qualitative match to the predicted changes provides support for RF-weighted normalization.

### Concluding Comments

While the intensity weighted normalization model we propose captures the vast majority of response variance in MT, several important questions remain. Notably, when measuring contrast response functions at different RF locations, our model does not fully explain the observed changes in response gain *R*_*max*_ (Figure 4). Additionally, although normalization has been implicated in mediating surround suppression,^1,46-48^ our model assigns a weight of zero to locations outside the classical RF, precluding their influence and potentially underestimating the full spatial extent of normalization. These observations suggest that intensity-weighted normalization, while powerful, may not capture all contributing sources of normalization.

Future research should investigate how intensity is encoded in higher-order cortical areas, where RF weight may not be spatially defined. By extending the intensity-weighted framework across cortical hierarchies, we may uncover deeper principles that govern how the brain balances excitation and suppression to construct stable, selective, and context-sensitive representations of the world.

## Materials and Methods

### Animal Preparation and Behavioral Task

All experimental procedures were approved by the Institutional Animal Care and Use Committee of the University of Chicago and followed the U.S. National Institute of Health guidelines. Under aseptic surgical conditions, two male monkeys (*Macaca mulatta*, 8.5 and 10.5 kgs) were implanted with a titanium head-post. Following a recovery period of four weeks, the monkeys were trained to do a visual change-detection task that controlled their attention. This task directs the animal’s attention away from the RFs of the neurons whose activity we recorded from to minimize attention-related modulations.

The animal initiated each trial by fixating on a small point that appeared on a calibrated CRT monitor (45° x 35°, 1024 × 768 pixels, 85 Hz frame rate, 57 cm viewing distance) against a gray background for 200 ± 35 ms. While maintaining fixation within a window (2.2-3.0° diameter for monkey A, 2.0-2.2° diameter for monkey M), the animals covertly attend to the left visual hemifield (ipsilateral to the recorded neurons) where a sequence of drifting Gabor stimuli appeared. Each Gabor appeared for 200 ms and was separated from the next by a variable interval of 200-400 ms. All Gabors shared the same orientation but drifted in one of two possible directions perpendicular to that axis, randomly selected on each presentation. For Monkey A the task Gabor had a SD of 1.5-2.0°while for monkey M the task Gabors had a SD of 1°.

Monkeys were rewarded with a juice reward if they responded within 500 ms of the appearance of a Gabor at a random place in the sequence that had a different orientation than the preceding Gabors. The monkeys reported detection by shifting their gaze from the fixation point to the location of the Gabor. On 10-40% of the trials, there was no change, and the animals were rewarded for maintaining fixation for the length of the trial (∼2-3 s). The trial ended early and without reward if the animal broke fixation.

The orientation of the task Gabor varied across sessions but was typically selected to differ from the recorded neuron’s preferred or non-preferred direction, as determined by mapping the neuron of interest’s direction tuning (see Neuronal Recordings). This design minimized the potential influences of spatial- or feature-based attention on neuronal responses.^49^

### Receptive Field (RF) Stimuli

While the monkeys performed the behavioral task, we presented full-contrast Gabor stimuli, drifting in either the preferred or non-preferred direction of an isolated neuron of interest. The Gabors appeared with an odd-symmetric phase and drifted by an integral number of sine wave cycles. For normalization measurements, these stimuli were shown at five iso-eccentric locations within the RF, chosen to span one half of the RF. We used iso-eccentric offsets to avoid anisotropies seen across the RF when transects span different eccentricities. The size of the Gabor stimuli varied across recording sessions and was scaled with the eccentricity of the RF (0.25-1.17° SD for Monkey A; 0.15-0.72° SD for Monkey M). The sequence of RF Gabors was presented with a fixed duty cycle (247 ms on, 200 ms off). In total we presented 31 single and paired stimulus configurations in the RF to measure normalization and the RF profile (see below).

Contrast response functions were measured separately, also by presenting Gabors in the RF while the monkeys performed the behavioral task. A single Gabor stimulus appeared at one of two RF locations and drifted either in the preferred or non-preferred direction. The size of the Gabor stimuli varied across recording sessions and was scaled for eccentricity (0.47-2.51° SD for Monkey A; 1.5° for monkey M), and assumed one of six contrast levels (3%, 6%, 12%, 25%, 50%, 100%). The 24 combinations of location, direction and contrast were used to construct four contrast response functions.

### Neuronal Recordings

Once each animal was proficient in the change-detection task, a recording chamber was implanted on the left hemisphere to allow for a posterior approach to MT (∼40-50° from horizontal in a parasagittal plane). Neuronal activity in MT was recorded using a linear multielectrode array (32 channel S-Probe; Plexon Inc.). On each recording day, we penetrated the dura with a guide tube that was targeted through a grid system.^50^ The electrode was then fed through the guide tube and advanced into cortex using a microdrive system (Kopf Instruments).

After isolating neurons in MT, we estimated the approximate RF location and preferred direction of one neuron using a hand-controlled visual stimulus. We then used computer-controlled Gabor stimuli to generate a direction tuning curve (12 directions) and map the RF using a 5×5 grid of azimuths and elevations using a small Gabor drifting in the preferred direction. The initial receptive field location mapping was followed by a finer mapping using a 5×5 grid localized around the RF center to further refine our estimation of the center.

Once the RF center was identified, we assigned a stimulus location there and added four additional locations along an iso-eccentric arc extending to one edge of the RF.^34^ The offset locations had uniform steps of chordal length from the center position. The assigned RF locations were used to measure how normalization varied across locations in the RF. We used Gabors drifting in either the preferred direction or one non-preferred direction. These directions were selected based on selected neurons direction tuning, with the non-preferred direction chosen to elicit approximately half the response of the preferred stimulus. To map the full width of the RF profile, randomly interleaved preferred-direction Gabors were presented individually at four iso-eccentric locations spanning the half of the RF not stimulated by the normalization stimuli.

We collected spikes using either the normalization or contrast RF stimuli while the monkey did the behavioral task. Recorded signals were amplified, bandpass filtered (250-7,500 Hz), and sampled at 30 kHz using a data acquisition system (Cerebus, Blackrock Microsystems). Both single and multiunits were recorded from 45 total sessions (28 from monkey A; 17 from monkey M). The continuous signal was spike sorted offline using an automated spike sorter (KiloSort 2.0 and KiloSort 4.0) and isolated units were curated using Phy.^51^

### Data Analysis

Data were analyzed using custom scripts written in MATLAB and Python. Only correctly completed trials were included in the analysis; trials with fixation breaks or early saccades were excluded. Spikes were counted during a window from 50 ms after stimulus onset to 50 ms after stimulus offset. To minimize variance associated with onset effects, we excluded the first stimulus in each trial’s stimulus train. To minimize attention-related response modulation, we excluded any stimuli that appeared with or after the target stimulus.

### Inclusion Criteria: Normalization Experiments

To be included for analysis, neurons were required to have activity statistically significant above the spontaneous activity to both the preferred and non-preferred stimuli at the RF center (one-sided Mann-Whitney U test, *α* = 0.05; Bonferroni-corrected where applicable). Additionally, the response to the preferred stimulus at the center had to be significantly greater than that to the non-preferred stimulus. Finally, the neuron had to show significant response to the preferred stimulus at one or more of the four peripheral locations.

For inclusion in the contrast response function analysis, neurons were further required to show significant responses at a minimum of two contrast levels for each of the following four conditions: preferred and non-preferred directions at both the RF center and the peripheral location. Additionally, we required that the Naka-Rushton function fit the data for all four conditions with a 95% confidence interval for the value of the *σ* parameter (obtained via bootstrapping) spanned less than a 30-fold range.

### Curve Fitting

For each unit that passed the inclusion criterion, we fit a suite of models to the mean firing rate across stimulus conditions using a non-linear least squares optimization (scipy.optimize.curve_fit). In the normalization experiments, we applied this method to fit a range of candidate models (Equations 1-7), with all parameters left unbounded. For the contrast response function experiments, we modelled responses using the Naka-Rushton function, defined as:

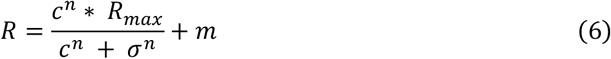

Here, *R* is the predicted response of the neuron, *c* is the stimulus contrast, *R*_*max*_ the maximum (saturated) response, *m* is the spontaneous rate, *σ* is the semi-saturation constant (i.e. the contrast at which the response is halfway between *R*_*max*_ and *m*), and *n* is an exponent determining the steepness of the curve. Parameters were constrained to be greater than zero.

### Gaussian RF profile

To quantify the spatial profile of each neuron’s RF, we fit the measured responses across RF locations to a one-dimensional Gaussian:

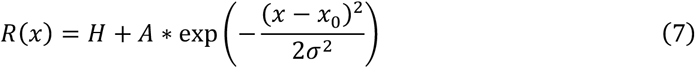

Where *R*(*x*) is the predicted response at spatial location *x, H* is a constant offset corresponding to the baseline activity, *A*, is the amplitude of the response above baseline, *x*_0_ is the location of the RF center, and *σ* is the standard deviation of the Gaussian.

### Bootstrap Estimation

As a part of the inclusion criteria for the contrast response function experiments, we employed a bootstrapping procedure to ensure that only neurons with reliable contrast sensitivity estimates were included in the analysis. Specifically, we estimated variability in the semi-saturation constant *c*_50_ for each of the four stimulus conditions by resampling trail-by-trial firing rates with replacement across 500 bootstrap iterations.

For each bootstrap sample, the Naka-Rushton function was re-fit to the resampled data, and the *c*_50_parameter was extracted. We then computed 95% confidence intervals for *c*_50_ by taking the 2.5^th^ and 97.5^th^ percentiles of the bootstrap distributions.

### Explainable Variance Estimation

To assess the fraction of neuronal response variability that our model could be expected to explain, we computed the explainable variance for each neuron using a two-step simulation-based procedure. This measure reflects the variance that can be accounted for by a noise-free version of the fitted model under the assumption of Poisson spiking variability.

First, we recovered the raw firing rates for each neuron by rescaling the normalized responses for all stimulus conditions. We then fit the intensity weighted normalization model to the raw responses using a non-linear least square procedure. Using the model parameters, we predicted the rates for each condition and generated simulated spike counts by drawing from a Poisson distribution, scaled for our 247 ms stimulus duration.

The number of simulated trials matched the number of trials for that neuron for each day.

We averaged the simulated spike counts per condition and refit the same model to these averaged (denoised) simulated responses. The explainable variance was computed as the coefficient of determination (R^2^) between the model predictions and the denoised simulated responses:

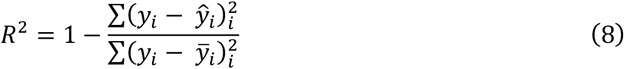

Where y_*i*_ are the simulated mean responses, *ŷ*_*i*_ are the model predictions based on the refitted parameters, and 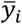 is the average of the simulated means.

We report the median explainable variance across the population. Finally, for each neuron, we recalculated the ratio of explained variance to the estimated explainable variance, using the explained variance score from the original model fits divided by the explainable variance. We summarize the distribution of these ratios using the median and interquartile range (IQR).

### Akaike Information Criterion (AIC)

We used the AIC to compare model performances when accounting for different numbers of parameters. Following model fitting for individual neurons, we computed the predicted responses and residual sum of square errors (RSS) and then the AIC was calculated for each model using the formula:

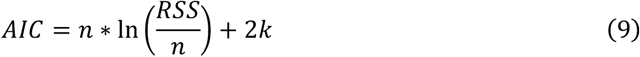

where *n* is the number of data points and *k* is the number of model parameters.

## Data availability

Data and code are available from the corresponding author upon reasonable request. All behavioral and neuronal data analysis were done using Matlab (MathWorks Inc.) and Python. Behavioral task was controlled using custom-written software (https://github.com/MaunsellLab/Lablib-Public-05-July-2016.git).

## Acknowledgements

We thank Jason N. Maclean, Jorge Jaramillo, Supriya Ghosh, Matthew C. Rosen, and Lai Wei for critical feedback on the manuscript; Rachel Parker and Autumn Mitchell for technical help with monkey procedures.

## Funding

This work was supported by National Institutes of Health grant, NIH RF1NS121772 (JHRM) and the Pritzker Fellowship from the Biological Sciences Division at the University of Chicago (CC). The funders had no role in study design, data collection and interpretation, or the decision to submit the work for publication.

## Author contributions

C.C. and J.H.R.M designed the experiments, performed the surgeries, and wrote the paper. C.C. performed the experiments and analyzed the data.

## Competing interests

The authors declare no competing interests.

## Declaration of generative AI and AI-assisted technologies in the writing process

During the preparation of this work the authors used ChatGPT (OpenAI) in order to assist with aspects of data analysis and manuscript preparation. All code related to experimental design, spike sorting, data organization, and assignment of spike counts to the appropriate stimulus condition arrays were performed by the authors. ChatGPT was primarily used to aid in generating code for figure formatting and visualization. In some instances, ChatGPT was also used to assist in generating code for fitting models to neuronal response data. All code was reviewed, validated, and modified by the authors to ensure accuracy and appropriateness for the analysis. ChatGPT was not used to draw scientific conclusions or interpret results. For manuscript preparation, ChatGPT was used solely to edit author-written text for clarity. It was not used to generate scientific content, interpret data, or describe figures. After using ChatGPT, the authors reviewed and edited the content as needed and take full responsibility for the content of the publication.

## Notes

### Competing Interest Statement

The authors have declared no competing interest.

